# Silver nanoantibiotics display strong antifungal activity against the emergent multidrug-resistant yeast *Candida auris* under both planktonic and biofilm growing conditions

**DOI:** 10.1101/2020.05.31.126433

**Authors:** Roberto Vazquez-Munoz, Fernando D. Lopez, Jose Lopez-Ribot

**Affiliations:** South Texas Center for Emerging Infectious Diseases, Department of Biology, The University of Texas at San Antonio, San Antonio, TX 78249, USA; The University of Texas at Austin, Austin, TX, 78712, USA

**Keywords:** Nanoantibiotics, Candida auris, Biofilms, Metallic nanoparticles, silver nanoparticles, antimicrobial nanomaterials

## Abstract

*Candida auris* is an emergent multidrug-resistant pathogenic yeast with an unprecedented ability for a fungal organism to easily spread between patients in clinical settings, leading to major outbreaks in healthcare facilities. The formation of biofilms by *C. auris* contributes to infection and its environmental persistence. Most antifungals and sanitizing procedures are not effective against *C. auris,* but antimicrobial nanomaterials could represent a viable alternative to combat the infections caused by this emerging pathogen. We have previously described an easy and inexpensive method to synthesize silver nanoparticles (AgNPs) in non-specialized laboratories. Here we have assessed the antimicrobial activity of the resulting AgNPs on *C. auris* planktonic and biofilm growth phases. AgNPs displayed a strong antimicrobial activity against all the stages of all *C. auris* strains tested, representative of four different clades. Under planktonic conditions, MIC values of AgNPs against the different strains were <0.5 μg mL^-1^; whereas calculated IC50 values for inhibition of biofilms formation were < 2 μg mL^-1^ for all but one of the *C. auris* strains tested. AgNPs were also active against preformed biofilms formed by all different *C. auris* strains, with IC50 values ranging from 1.2 to 6.2 μg mL^-1^. Overall, our results indicate potent activity of AgNPs against strains of *C. auris*, both under planktonic and biofilm growing conditions, and indicate that AgNPs may contribute to the control of infections caused by this emerging nosocomial threat.

## Introduction

*Candida auris* is an emergent multidrug-resistant yeast that has been reported worldwide since its detection in Japan in 2009 [1]. It has been determined that there are four geographic clades of this pathogen, including South Asian (clade I), East Asian (clade II), South African (clade III) and South American (clade IV), which interestingly seemed to have emerged independently in different regions of the world at the same time [2,3], with a potential fifth clade recently identified in Iran [4]. *C. auris* is described as an ovoid-shaped non-dimorphic yeast that rarely forms pseudohyphae and exhibits two growing typical phenotypes: aggregative and non-aggregative [5,6]. *C. auris* spreads in healthcare settings, posing a risk for hospital patients due to its high mortality rate invasive infections, and its healthcare-associated outbreaks [6,7]. It easily contaminates surfaces and medical instrumentation within healthcare facilities for long periods, which poses a risk factor in healthcare facilities worldwide [7]. *C. auris* is considered as an urgent threat by the Centers for Disease Control and Prevention, according to their “Antibiotic Resistance Threats in the United States, 2019” [8].

Currently, the prevention and treatment of *C. auris* are challenging due to several factors. This yeast is known for its resistance to the main classes of clinically-used antifungal agents, and it is usually found as resistant to multiple drugs; also, its antifungal-resistance profile is different in each strain [9], which negatively impact treatments effectivity. Additionally, it is commonly misidentified in clinical laboratories, often leading to inappropriate treatments. Furthermore, it is able to form biofilms, and *C. auris* biofilms, besides being intrinsically resistant to all antifungal agents [10], can also withstand exposure to harsh setting conditions, such as high temperature and salinity concentration, and can survive in plastic surfaces up to for two weeks [11]. This yeast is highly resistant to current sanitation processes and treatments, such as ultraviolet light and quaternary ammonium compounds [5], which defy our capacity to control its propagation.

Therefore, new treatments are needed to prevent and control *C. auris* growth and dissemination. Nanotechnology can provide new cost-effective antimicrobial nanomaterials (nanoantibiotics) that work as disinfectants and antimicrobial drugs. In particular, silver nanoparticles (AgNPs) exhibit good antimicrobial properties with a wide range of action against a broad range of microorganisms, including several Candida species (Raghunath & Perumal, 2017; Vazquez-Muñoz et al., 2017). Additionally, nanoantibiotics can overcome the microbial drug-resistance to antibiotics (Rudramurthy et al., 2016; Vazquez-Muñoz et al., 2019). However, to date, only one report from our group has described the effect of AgNPs (synthesized using a different method) against a single isolate of *C. auris* in suspension and on functionalized medical and environmental surfaces [16]. This study demonstrated that silver nanoparticles effectively inhibit the *C. auris* biofilm formation. Additionally, a non-nanostructured silver commercial formulation (0.01% silver nitrate with 11% hydrogen peroxide) was shown to be effective against *C. auris* (Biswal et al., 2017).

We have recently reported on a modified facile, inexpensive synthetic method to generate silver nanoparticles (AgNPs) in non-specialized laboratories, and described their antibacterial and antifungal properties. We hypothesize that AgNPs display strong antifungal activity against multiple strains of *C. auris,* regardless of their clade, antibiotic-resistant profile, or morphological traits. Therefore, the objective of this study was to assess the antimicrobial activity of AgNPs synthesized using our newly described method, on different *C. auris* strains, for which we have evaluated the antimicrobial activity of nanoantibiotics against ten *C. auris* strains from the CDC panel, representing the four different major clades, both under planktonic and biofilm growing conditions.

## Materials and Methods

### Reagents

Roswell Park Memorial Institute (RPMI) 1640 culture medium, Phosphate Saline Buffer (PBS), 2,3-Bis(2-methoxy-4-nitro-5-sulfophenyl)-2H-tetrazolium-5-carboxanilide salt (XTT), and Menadione, were acquired from Sigma-Aldrich (MO). Osmium Tetroxide (OsO4) (4%) and Glutaraldehyde from Ted Pella Inc. Solutions of the different reagents were prepared in Milli Q water.

### Silver nanoparticles

Silver nanoparticles (AgNPs) coated with polyvinylpyrrolidone (PVP) were synthesized by a chemical reduction protocol, reported previously by our group (Vazquez-Muñoz et al., 2019). The synthesis method uses a simple and fast chemical reduction process that involves the addition of a PVP to a warmed silver nitrate solution, followed by sodium borohydride. The AgNPs obtained have an aspect ratio close to 1, an average size of 6.18 ± 5 nm, and a zeta potential score of −16.2 mV. The negative-charged, small, spheroid AgNPs displayed strong antimicrobial activity against *S. aureus* and *C. albicans* (Vazquez-Muñoz et al., 2019). This easy-to-replicate-synthesis method was specifically developed so that it can be readily implemented in non-specialized facilities and laboratories.

### Strains and culture conditions

*Candida auris* strains were acquired from the Centers for Disease Control and Prevention (CDC) Antimicrobial Resistance (AR) Isolate Bank stock [9].The following AR Bank strains were used: Clade I (#0382, #0387, #0388, #0389, and #0390), Clade II (#0381), Clade III (#0383 and #0384), and Clade IV (#0385, #0386). Frozen glycerol stocks of the microbial cells were subcultured onto Yeast extract-Peptone-Dextrose (YPD) (BD Difco, MD) agar plates, for 48 h at 37 °C. Then, *C. auris* was cultured into YPD liquid media overnight at 30 °C, in an orbital shaker. Cells from these cultures were prepared for the susceptibility tests, as described in the following sections.

### Antifungal susceptibility testing under planktonic conditions

The antimicrobial activity of the nanoparticles on the *C. auris* planktonic cells was determined following the guidelines from the CLSI M27 protocol [18] for *Candida* species, with minor modifications. Briefly, the yeast cells were washed twice in PBS, counted in a Neubauer chamber, and adjusted for a final concentration of 10^3^ cells mL^-1^ in RPMI culture media. Then, 50 μL of the *C. auris* strains were inoculated in 96 multi-well round-bottom plates (Corning Inc., Corning, NY). AgNPs were prepared in a two-fold dilution series in RPMI, then 50 μL of the dilution series were added to the plates with the yeast, for final AgNPs concentration range from 0.5 to 256 μg mL^-1^. Plates were incubated at 35 °C for 48 h. The Minimal Inhibitory Concentration (MIC) was set as the concentration in the well at which no microbial growth was observed. The Minimal Fungicidal Concentration (MFC) was also established, as follows: after reading the MIC in each plate, 10 μl from each well containing the untreated and treated microbial cells were re-inoculated in YPD agar plates and incubated for 24 h at 37 °C. The MFC was set as the lowest concentration of nanoparticles that displayed the growth of three or fewer colony-forming units (CFUs). To ensure reproducibility, the experiment was independently performed by two people, on separate days, using different batches of AgNPs and *C. auris* cultures. Experiments were performed using duplicates of the plates which contained triplicates of each condition.

### Antibiofilm activity assays

The antibiofilm activity of AgNPs was evaluated in both the biofilm formation phase and on the preformed biofilm as previously reported by our group [19]. For inhibition of biofilm formation overnight cultures of *C. auris* yeast cells were washed twice in PBS and adjusted to 2×10^6^ cells mL^-1^ in RPMI culture media. Fifty μL of the adjusted cell suspension were transferred to flat-bottom 96-multiwell plates (Corning). Then, 50 μL of AgNPs prepared in a two-fold dilution series were added into multi-well plates, for a final concentration range from 0.5 to 256 μg mL^-1^, with appropriate positive and negative controls. The plates were then incubated at 37 °C for 24 h to allow for biofilm formation. We also tested the activity against preformed biofilms. Briefly, cells from overnight liquid cultures were washed twice in PBS and adjusted to 1×10^6^ cells mL^-1^ in RPMI. Then, 100 μL of the microbial suspension were inoculated into 96-multiwell plates and then incubated for 24 h at 37 °C. After incubation, the preformed biofilms were washed twice in PBS. Then, 100 μL of AgNPs in two-fold dilutions series (prepared in RPMI medium and resulting in final concentrations ranging from 512 to 1 μL mL^-1^) were transferred to the wells of the microtiter plates with the preformed biofilms. Finally, the plates were incubated at 37 °C for an additional 24 h.

The AgNPs anti-biofilm activity was determined using the XTT colorimetric method [19] for both inhibition of biofilm formation and the preformed biofilm stages. Briefly, at the end of the procedure biofilms were washed twice with PBS, then 100 μL XTT/menadione was added to each well containing treated and untreated biofilms, and in the empty wells (blank). Plates were protected from light and incubated at 37 °C for 2 hours. XTT absorbance was measured at λ=490 nm in a Benchmark Microplate Reader (Bio-Rad Inc). From the collected data, we generated dose-response curves to assess the IC50 – the drug concentration required to reduce the biofilm activity by 50%-, by fitting the normalized results to the variable slope Hill equation (for assessing the nonlinear dose-response relationship) using Prism 8 (GraphPad Software Inc). To verify the reproducibility of the antibiofilm activity, the experiment was independently repeated by two different people. AgNPs from different rounds of syntheses were tested using two replicates of multi-well plates, each with three replicates of the treatments.

### Ultrastructural analysis

We assessed the effect of AgNPs on the biofilm structure in all *C. auris* strains from the four clades, using the biofilm inhibition assays. The biofilms were treated with sub-lethal (yet still inhibitory) concentrations of AgNPs. Treated and non-treated (control) biofilms structural analysis was performed using optical and scanning electron microscopy. Biofilms were washed twice with PBS, then fixed with a 2.5 % glutaraldehyde solution for 3 h at 4 °C. For the Optical Microscopy observations, the glutaraldehyde-fixed biofilms were observed under a 400x magnification using the bright field mode, in an inverted optical microscope (Fisher Scientific). For the Scanning Electron Microscopy analysis, the glutaraldehyde-fixed biofilms were post-fixed and stained with a 1% Osmium tetroxide solution, for 2.5 h at 4°C. Then, the biofilms were dehydrated in an ascending concentration ethanol series, from 30% to 100%. Finally, the ethanol was completely removed, and the dried samples were coated with gold, with 25 milliamperes current for 3 minutes, in a Sputter Coater SC7620 (Quorum Technologies). The gold-coated biofilms were observed in a TM4000Plus Scanning Electron Microscope (Hitachi Inc.), with a magnification 500 and 2500x, at a voltage of 10 KeV in the High Vacuum mode. The samples were prepared in duplicates and different fields of both replicates from each sample were observed.

## Results and Discussion

### Silver nanoparticles inhibit the planktonic growth of *C. auris*

AgNPs exerted a strong antimicrobial activity against all the *C. auris* strains growing under planktonic conditions. Against 9 out of 10 strains, AgNPs MIC values were < 0.5 μg mL^-1^, and the MFC values were only slightly higher, ranging from 1 to 2 μg mL^-1^ for all strains, except for AR #0381 which had an elevated MFC of 32 μg mL^-1^. AgNPs MIC and MFC values against each strain are summarized in **Table 1**. Our results show that AgNPs display potent antifungal activity at very low concentrations in virtually all *C. auris* strains tested. In a previous report (Vazquez-Muñoz et al., 2019), similarly synthesized AgNPs exhibited strong antibacterial and antifungal activity. The MIC for *S. aureus was* 4 μg mL^-1^, whereas for *C. albicans* the MIC was 2 μg mL^-1^, which is similar to the most common antifungals. It is worth noting that all *C. auris* strains are more susceptible to the AgNPs than *C. albicans* tested under similar experimental conditions. Also, the anti-candidal activity of these AgNPs against planktonic cells of *C. auris* parallels other studies using different nanoparticles against different *Candida* species with typical MIC values in the 1 – 10 μg mL^-1^ range (Monteiro et al., 2011; Patra & Baek, 2017; Vazquez-Muñoz et al., 2017; Wady et al., 2014). Moreover, all *C. auris* strains tested here displayed susceptibility to AgNPs, irrespective of their growth characteristics, susceptibility profiles against conventional antifungal, or their geographical origin (clade). This has been observed in other microorganisms, including other Candida species, where the drug-resistant strains and drug-sensitive strains from the same species display a similar susceptibility (MIC value) to AgNPs [23,24]. Additionally, our results suggest that AgNPs antimicrobial performance is better than the main antifungals on all the tested *C. auris* CDC AR strains, according to their antifungal susceptibility profile reported by the CDC [25]. Although there are not established MIC breakpoints for the main available antifungals against *C. auris*, the tentative MIC Breakpoints for some of them are the following: fluconazole >32 μg mL^-1^, amphotericin B >2 μg mL^-1^, caspofungin >2 μg mL^-1^, and micafungin >4 μg mL^-1^ [25]. The AgNPs MIC (<0.5 μg mL^-1^) outperforms even the most potent antifungal drug.

**Table 1.**
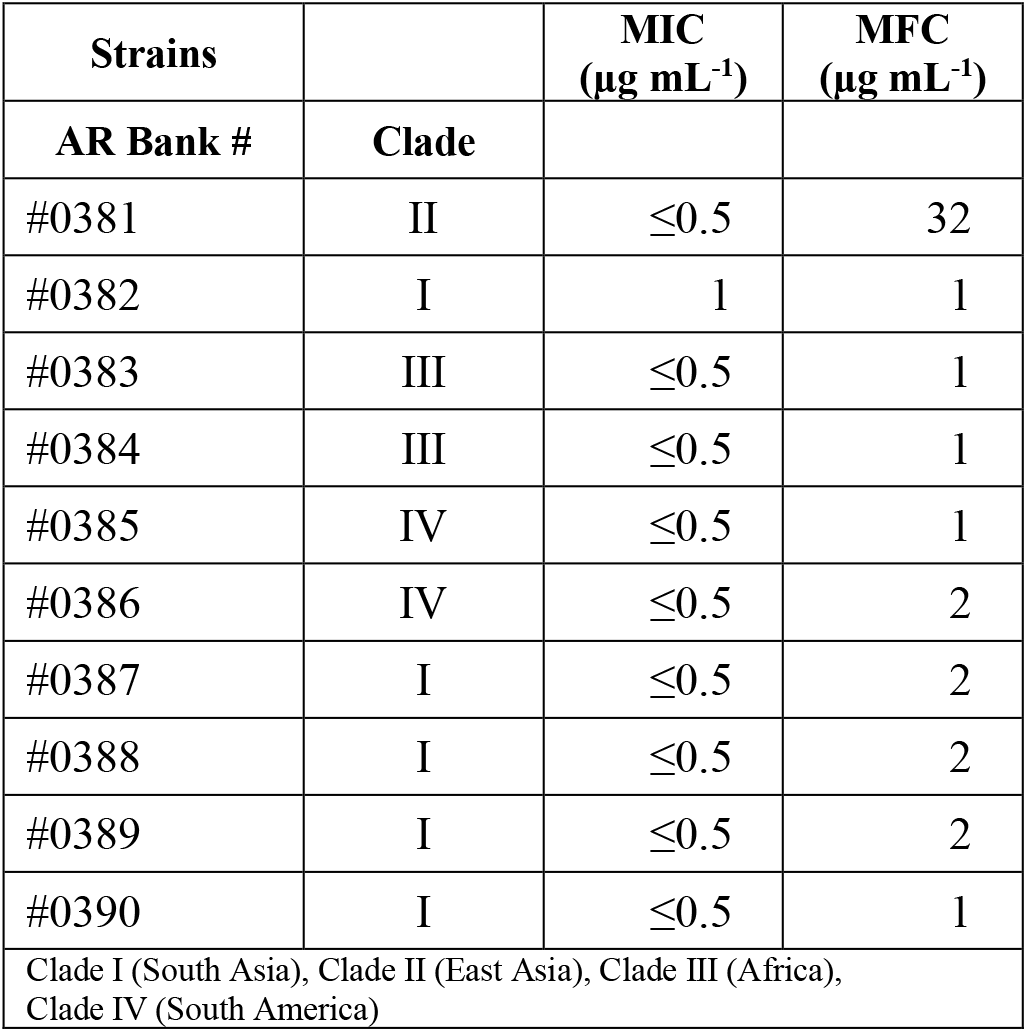
MIC/MFC values of AgNPs against *C. auris strains* under planktonic growth conditions.

| Strains | | MIC<br>($\mu\text{g mL}^{-1}$) | MFC<br>($\mu\text{g mL}^{-1}$) |
| --- | --- | --- | --- |
| AR Bank # | Clade |  |  |
| #0381 | II | $\leq 0.5$ | 32 |
| #0382 | I | 1 | 1 |
| #0383 | III | $\leq 0.5$ | 1 |
| #0384 | III | $\leq 0.5$ | 1 |
| #0385 | IV | $\leq 0.5$ | 1 |
| #0386 | IV | $\leq 0.5$ | 2 |
| #0387 | I | $\leq 0.5$ | 2 |
| #0388 | I | $\leq 0.5$ | 2 |
| #0389 | I | $\leq 0.5$ | 2 |
| #0390 | I | $\leq 0.5$ | 1 |
| Clade I (South Asia), Clade II (East Asia), Clade III (Africa),<br>Clade IV (South America) |  |  |  |

This may be due to the proposed mechanisms of action of AgNPs. The mechanism of action of the antifungal drugs is linked to specific molecular targets that disrupt the cell metabolism or structure, affecting growth. In response to these stresses, specific but relatively small changes at the structural or molecular level may increase their probability to resist the action of antifungals, as previously described for *C. auris* [26,27]. In contrasts, AgNPs cause several simultaneous types of structural and metabolic damage in the *Candida* cells, such as membrane depolarization [28], cell wall/membrane disruption [29], increase in ROS production [30], inhibit enzymatic function [31], cell arrest [28], among many others. This massive disruption of cellular structure and function reduces their ability to withstand the AgNPs effects. Therefore, the metabolic/structural differences among the different strains are not significant when the cells from different *C. auris* strains are exposed to AgNPs.

### Silver nanoparticles inhibit *C. auris* biofilm formation

*C. auris* is capable of forming biofilms that improve their adherence to surfaces [6] and increase their resistance to antifungal drugs [5,10]. The mechanisms that enhance their resistance are mostly unknown, but some factors are known to help *C. auris* withstand harsh conditions include the protection by matrix polysaccharides [32] and the overexpression of efflux pumps [33]. Thus, the formation of *C. auris* biofilms represents a current threat to both individual patients and healthcare facilities [7]. In previous work, we demonstrated the antimicrobial activity of the AgNPs on the planktonic stage of *C. albicans* (Vazquez-Muñoz et al., 2019), although their activity was not assessed on the biofilm stages. In this work, we evaluate the anti-biofilm activity of the AgNPs during the biofilm formation phase and against fully mature, preformed biofilms.

AgNPs exhibited a strong activity to prevent biofilm formation in *C. auris,* regardless of the clade. **Figure 1** shows the biofilm-inhibitory effect against representative isolates from each clade, including strains AR #0381 (clade I), #0383 (clade III), #0386 (clade IV), and #0390 (clade II). The AgNPs antibiofilm activity against all 10 strains tested is shown in supplementary **figure S1A** (supplementary materials). The calculated IC50 values ranged from 0.5 to 4.9 μg mL^-1^ (**Table 2**), and for 9 out of 10 strains, the IC50 values were < 2 μg mL^-1^. These results indicate that AgNPs exert a potent activity for the prevention of biofilm formation by the different *C. auris* strains. Interestingly, these values are only slightly higher than the MIC values obtained under planktonic growth. These values also compare favorably to those described before for conventional antifungals against biofilm formation for some of the same *C. auris* strains (Dekkerová et al., 2019). We note that the AgNPs IC50 value for the *C. auris* AR #0390 strain is higher than the value reported for the same strain as reported by Lara *et al* (1.1 μg mL^-1^ vs 0.06 μg mL^-1^, respectively), which is most likely related to the different techniques used for the synthesis of these nanoantibiotics resulting in AgNPs with different characteristics. Also, the antibiofilm activity of our AgNPs is comparable to the activity described for AgNPs synthesized using different methods against other *Candida* species [29,34,35].

**Figure 1.**
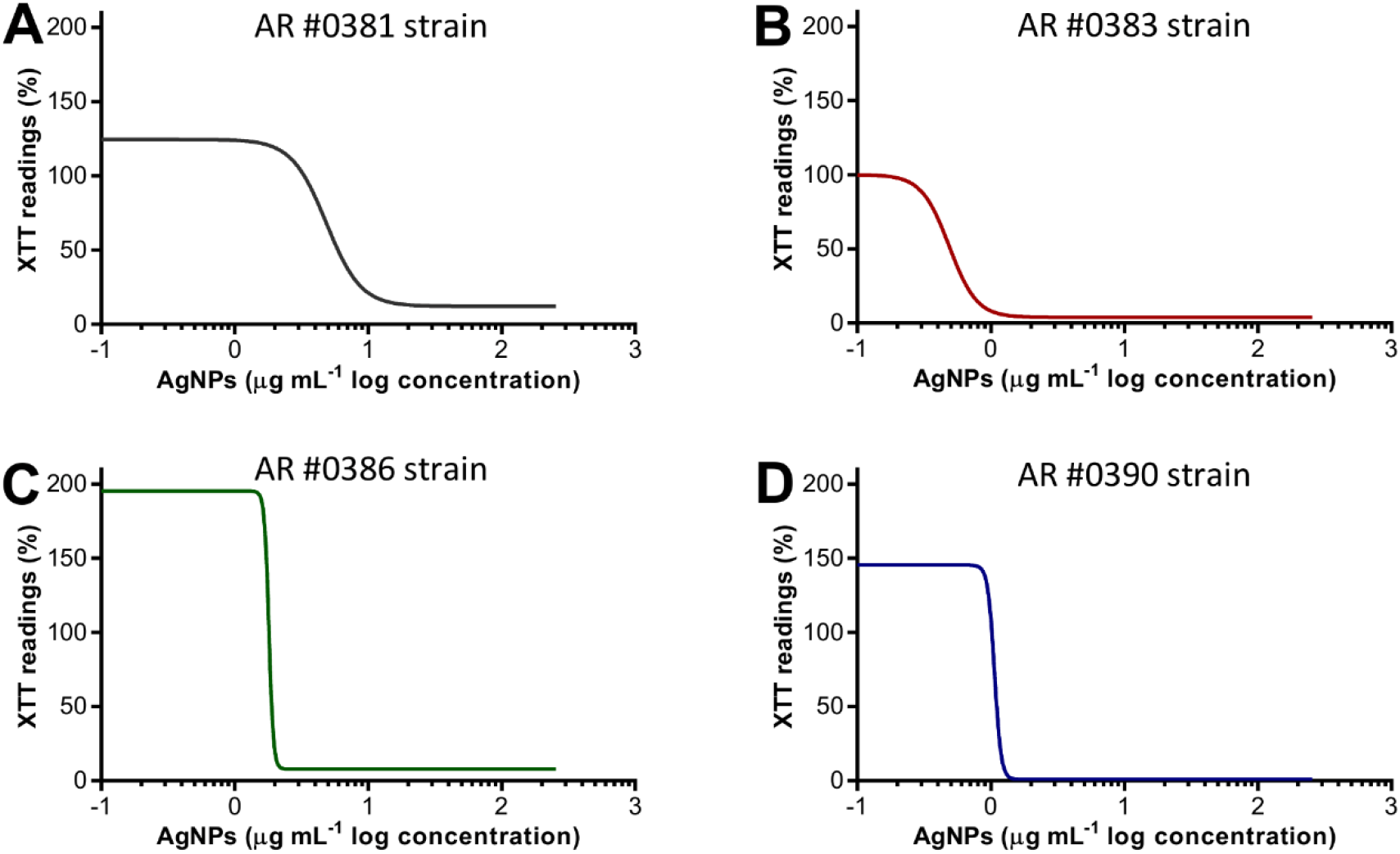
AgNPs inhibit the biofilm formation on *C. auris*. The dose-response curve shows that AgNPs display potent inhibitory activity, expressed as XTT readings against the *C. auris* AR #0381 (clade I), #0383 (clade III), #0386 (clade IV), and #0390 (clade II) strains during the biofilm formation phase.

**Table 2.**
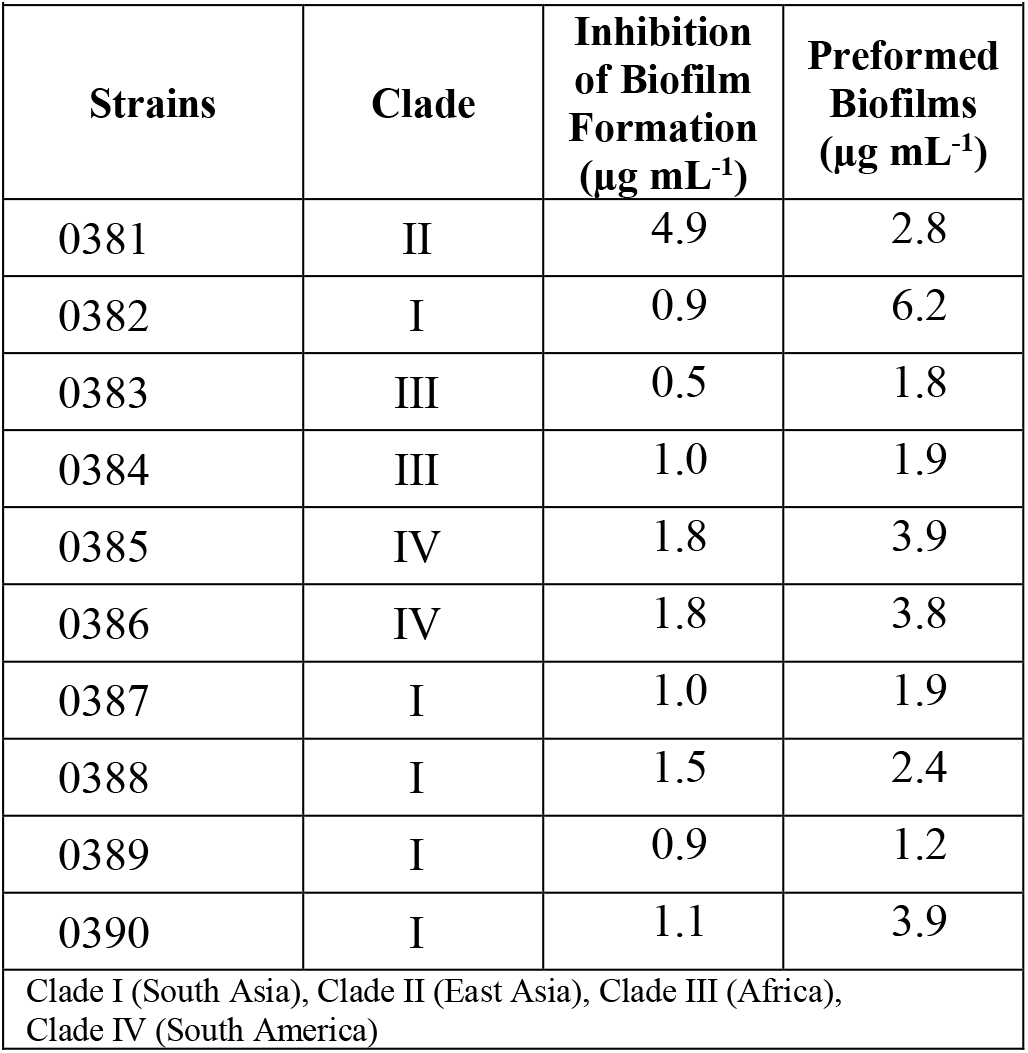
Calculated IC50 values for AgNPs against *C. auris* biofilms by the different strains.

Interestingly we observed that some strains exhibited a significant increase in the biofilm activity (determined by the XTT readings) when grown in the presence of very low concentrations of AgNPs. This effect was consistently observed in all replicates, although with different degrees of intensity. To the extent of our knowledge, this phenomenon has not been observed in yeasts treated with AgNPs, but it has been previously reported in bacteria [36]. Nevertheless, this increase in activity is promptly extinguished at just slightly higher concentrations of AgNPs. Additionally, to assess if the augmented activity was specific to the AgNPs, we evaluated the influence of silver nitrate on the *C. auris* AR #0390 strain, under the same culture conditions used for the AgNPs susceptibility assays. We observed an increase in the biofilm activity in subinhibitory concentrations of silver ions (**Fig S2,** supplementary materials). This effect on the biofilm activity must be addressed in further studies, to determine the potential disadvantage of low-silver-content products intended against *C. auris* biofilms. As mentioned above, there are commercially available products containing silver [37], and therefore their viability for combat the biofilms of microbial pathogens must be assessed.

### Silver nanoparticles display antibiofilm activity against preformed *C. auris* biofilms

It is well known that once a biofilm is established, candidal cells within these biofilms display increased susceptibility to most clinically-used antifungal agents [10,38]. This is particularly true in the case of *C. auris* biofilms, which are intrinsically resistant to all three main classes of antifungals (polyenes, azoles, and echinocandins) as well as to physical and chemical sanitizing methods [5,10]. As seen in **Fig. 2**, the AgNPs also displays potent activity against fully mature, preformed biofilms of *C. auris*, irrespective of their clade, as observed for representative isolates AR #0381 (clade I), #0383 (clade III), #0386 (clade IV), and #0390 (clade II). The AgNPs activity on the preformed biofilms from all 10 *C. auris* strains tested is shown in supplementary **figure s1B** (supplementary materials). From the dose-response experiments, the resulting calculated IC50 values of AgNPs against preformed biofilms of the different *C. auris* strains ranged from 1.2 to 6.2 μg mL^-1^ (**Table 2**), and were < 4 μg mL^-1^ for 9 out of the 10 strains. Interestingly, these values are similar (typically within one fold dilution) to those observed for the same strains in the case of biofilm inhibition (compare values in both columns of **Table 2**). Therefore, in stark contrast with conventional antifungal agents, the AgNPs potency does not seem to be particularly reduced after the biofilm has reached maturity. Moreover, the AgNPs antifungal activity against the preformed biofilms is equivalent or even better to that of conventional antifungals. For the *C. auris* AR strains #0383, #0386, and #0390, the AgNPs anti-biofilm activity is superior to the activity of Fluconazole (range from >64 to >1024 μg mL^-1^) and Caspofungin (>16 μg mL^-1^). Furthermore, the AgNPs potency parallels to the activity of Amphotericin B (1 to > 8 μg mL^-1^) [34,39]. Overall, although multiple mechanisms confer resistance of cells within biofilms against conventional antifungal agents [40], our results suggest that these do not equally affect the anti-biofilm activity of AgNPs. We note that similar to our observations in the biofilm-inhibitory assays described before, we also observed an increase in the biofilm activity (determined by the XTT readings) at very low AgNPs concentrations (**Fig 2** and Supplementary **Fig S2**).

**Figure 2.**
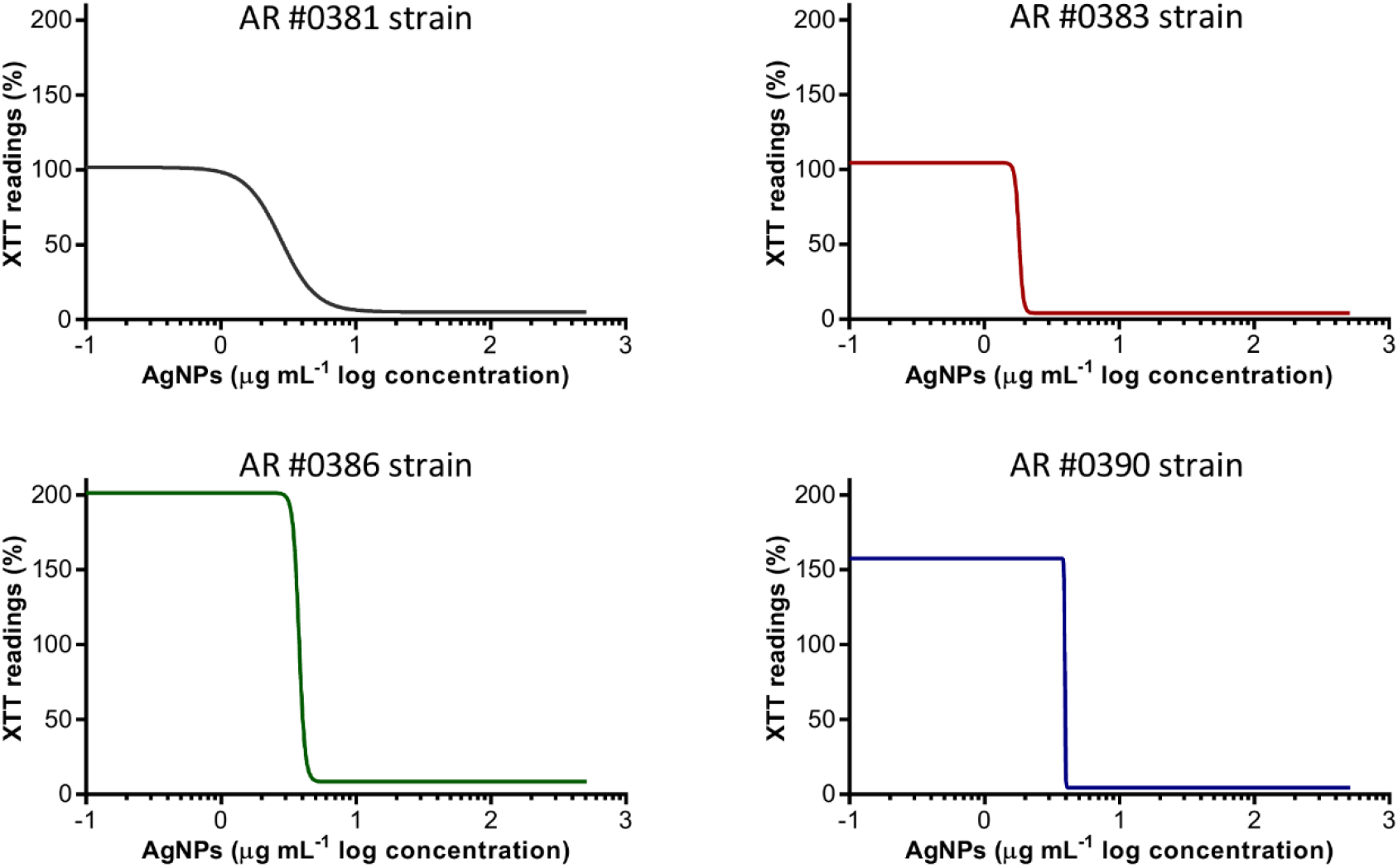
Antibiofilm activity of AgNPs against *C. auris* preformed biofilms. The dose-response curve shows that AgNPs display potent inhibitory activity-expressed as the XTT readings-against preformed biofilms of the *C. auris* AR #0381 (clade I), #0383 (clade III), #0386 (clade IV), and #0390 (clade II) strains.

### Alterations of *C. auris* biofilm structure due to the inhibitory activity of AgNPs

Once we had established the activity of AgNPs against *C. auris* biofilms, we were interested in the visualization of the effects of treatment with these nanoantibiotics exert on the overall biofilm structure, as well as on individual cells within the biofilms. Thus, in another set of experiments, we grew biofilms of the all different *C. auris* strains in the presence of subinhibitory concentrations of the AgNPs, with results for a representative strain from each clade shown in Figures 2, 3, 4, and 5, corresponding to strains #0381 (East Asia clade), #0383 (Africa clade), #0386 (South America clade), and #0390 (South Asia clade) respectively. Optical microscopy revealed that AgNPs decrease the ability of *C. auris* to form biofilms. As seen in supplementary **figure S3** (supplementary materials), in the untreated control samples, biofilms formed by the different strains uniformly covered most of the bottom of the wells in the microtiter plates. In contrast, inhibitory concentrations of AgNPs disrupt the biofilm formation in all *C. auris* strains, as revealed by the noticeable reduction of the coverage area of biofilms on the bottom of the wells. At higher concentrations of AgNPs, biofilm formation was drastically reduced, with only isolated cells scattered on the bottom of the wells being visible under the microscope.

**Figure 3.**
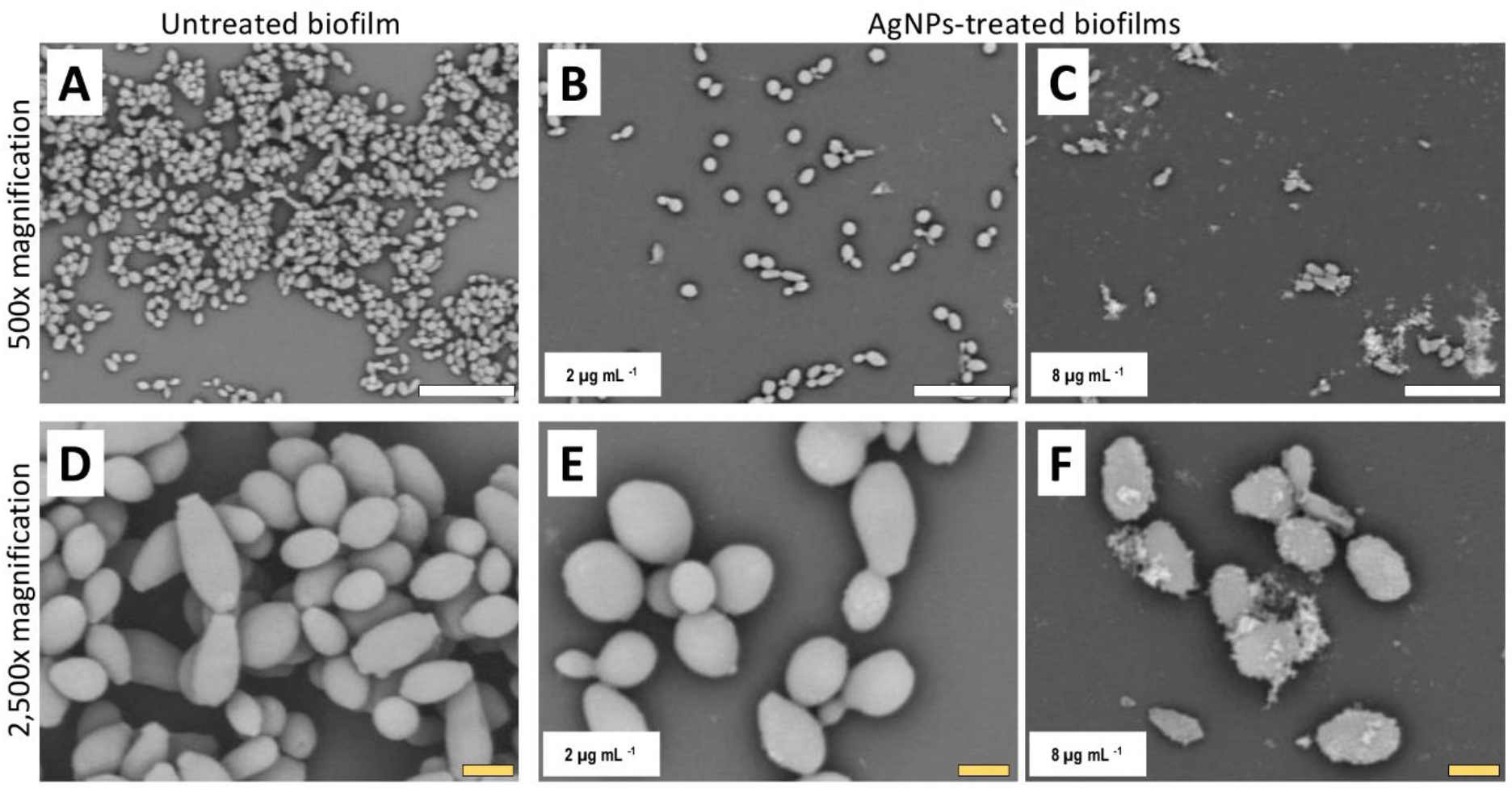
AgNPs affect biofilm structure and cellular morphology of *C. auris* strain #0381. SEM micrographs reveal that untreated biofilms have a larger area of distribution (**A**) than the AgNPs-treated biofilms (**B, C**). Also, the shape and size of the cells are affected by the AgNPs (**E, F**), whereas the untreated biofilms remain unaltered (**D**). Scale bar: white=20 μm, yellow=2 μm.

**Figure 4.**
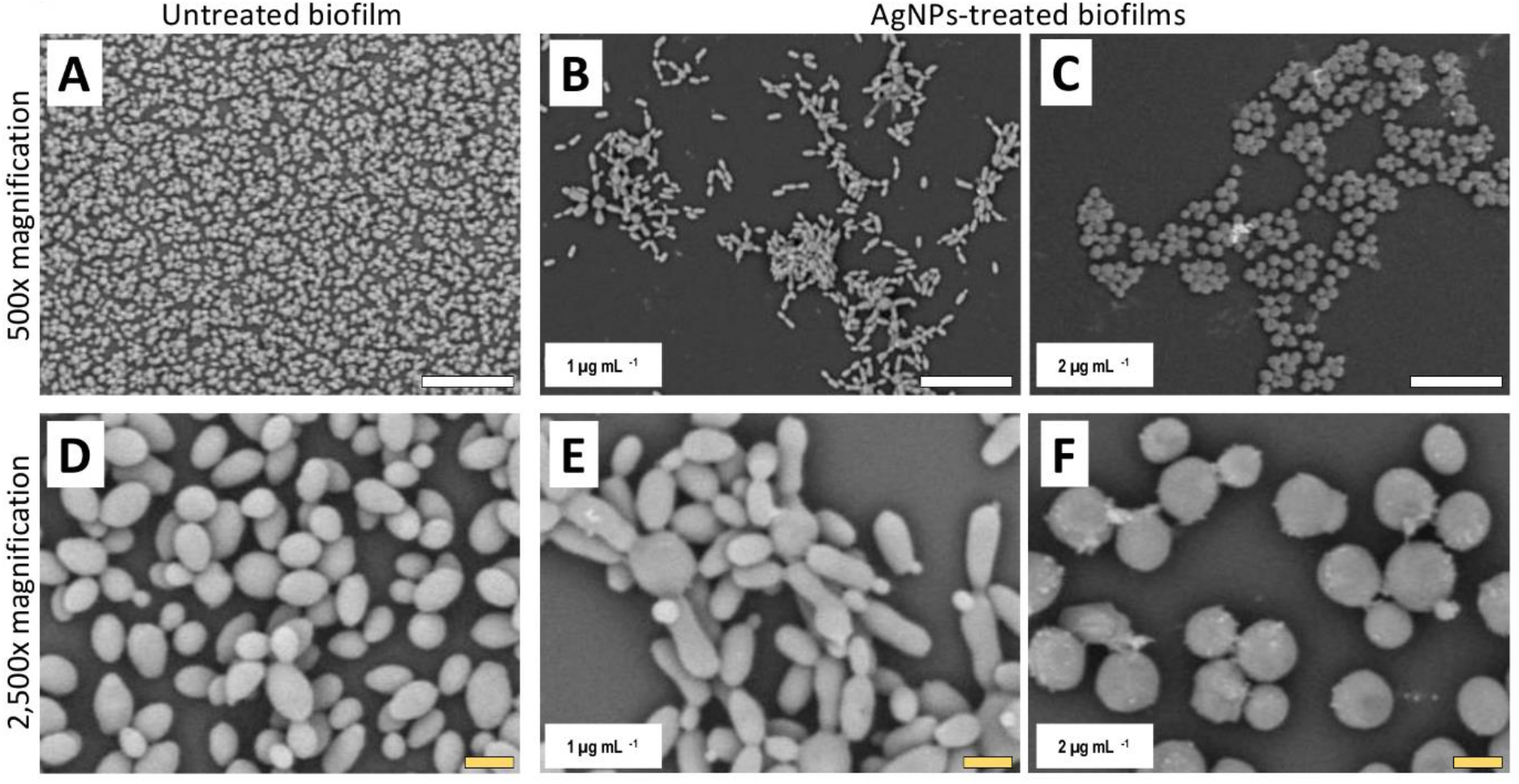
AgNPs reduce biofilm formation of *C. auris* strain #0383. SEM micrographs show a noticeable reduction in the biofilm formation in the AgNPs-treated biofilms (**B, C**) when contrasted with the untreated biofilms (**A**). However, the impact on cell morphology appears to be minimal (**E, F**), as the cell shape and size of cells within the treated biofilms are similar to those of the untreated control (**D**). Scale bar: white=20 μm, yellow=2 μm.

**Figure 5.**
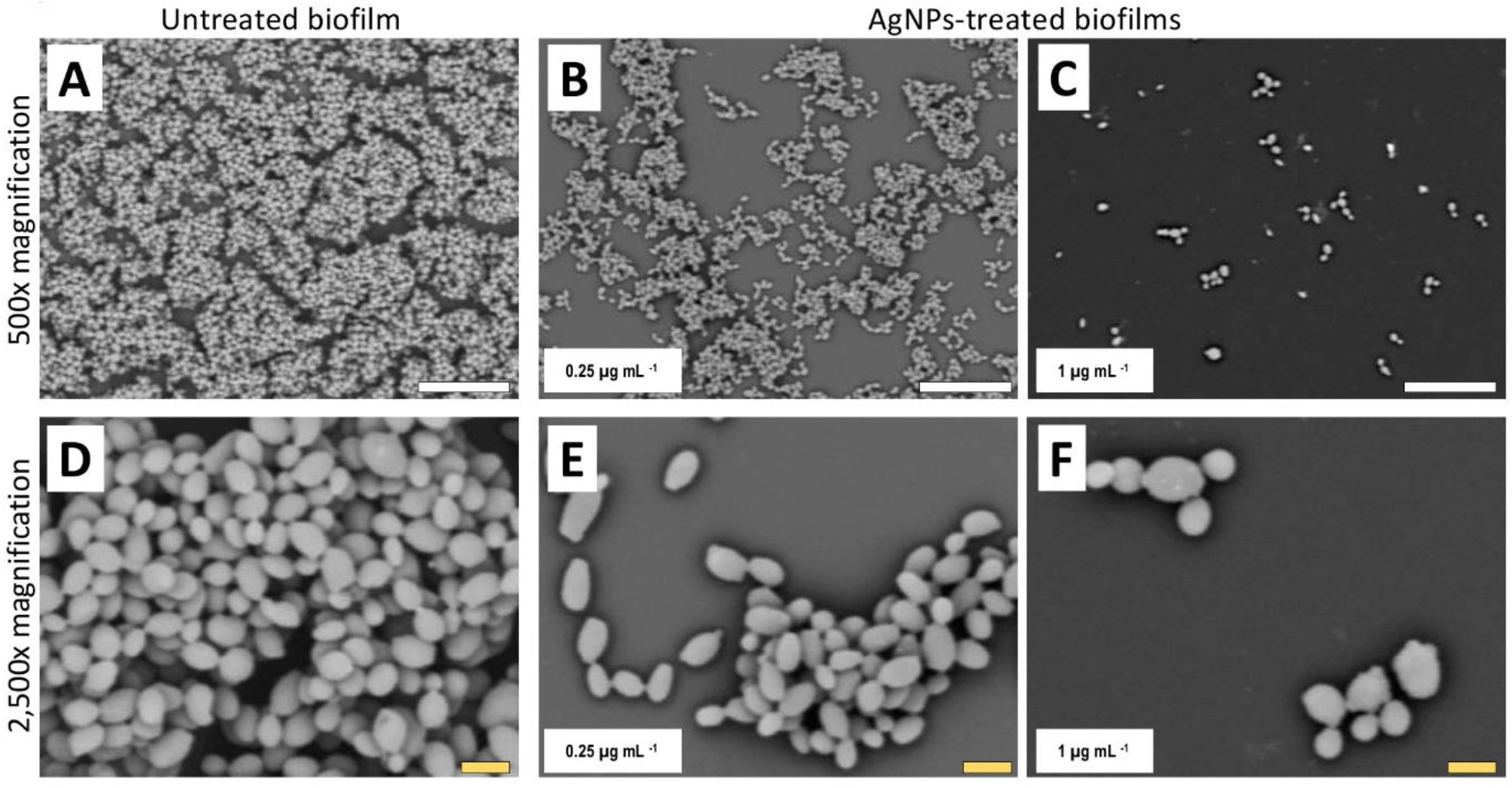
AgNPs affect biofilm structure and cellular morphology of *C. auris* strain #0386. SEM micrographs show that biofilms and individual cell morphology are drastically affected by the AgNPs. Treated biofilms (**B, C**) show an evident decrease in coverage area as compared to untreated biofilms (**A**), whereas the cell morphology is changed by treatment with AgNPs (**E, F**) as compared to cells in untreated biofilms (**D**). Scale bar: white=20 μm, yellow=2 μm.

The biofilms were observed using SEM at a low (500x) and a high (x2,500) magnifications, to further determine the effect of treatment with AgNPs on the biofilm structure and the cell morphology. *C. auris* from the distinct clades display differences in the cell morphology and the biofilm organization. SEM images confirmed that exposure to inhibitory concentrations of silver nanoparticles decreases the biofilm-forming ability of the different *C. auris* strains (**Figures 3** to **6** and **supplementary Figure S4**). SEM micrographs showed that untreated biofilms display a uniform distribution with a tight clustering of cells; in contrast, AgNPs-treated biofilms cover a noticeably lesser area and the cells appear to be less clustered. This finding is similar to that reported recently by Lara *et al,* for *C. auris* strain #0390 when exposed to a different type of AgNPs [16].

**Figure 6.**
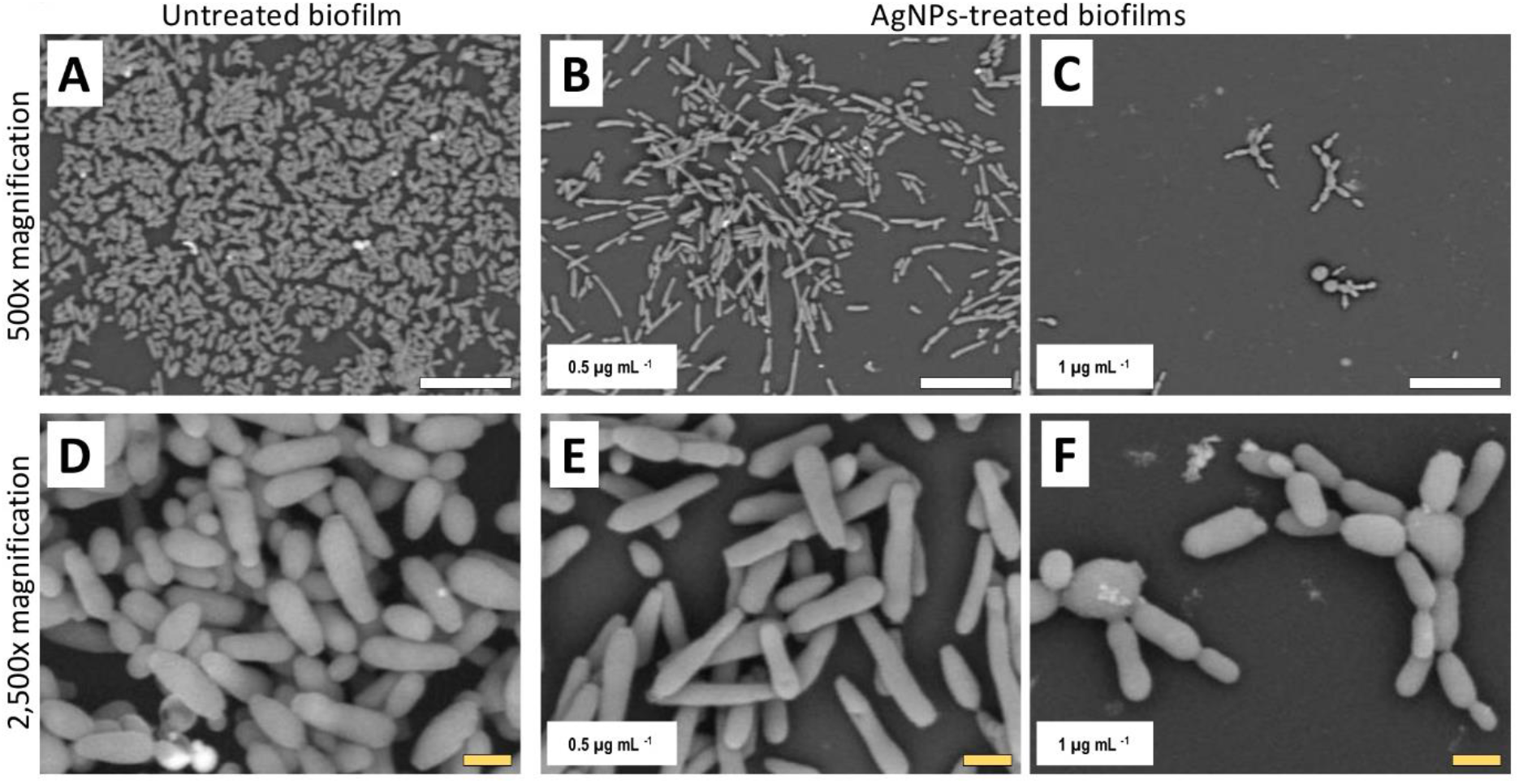
AgNPs affect biofilm structure and cellular morphology of *C. auris* strain #0390. The SEM micrographs of *C. auris* #0390 strain reveal that untreated biofilms (**A**) display a larger area of coverage than the AgNPs-treated biofilms (**B, C**). Moreover, untreated cells display pseudohyphae-like shape (D), which is also observed at the lowest concentration of AgNPs (**E**), but higher concentrations of AgNPs induce an aberrant morphology on cells and reduce the cell separation process (**F**) as compared to cells in untreated biofilms (**D**). Scale bar: white=20 μm, yellow=2 μm.

Moreover, when observed at higher magnification, it was revealed that treatment with AgNPs damages the fungal cell structure. In the control (untreated) samples, cells within the biofilms formed by strains #0381 (**Fig. 3**), #0383 (**Fig. 4**) and #0386 (**Fig. 5**) displayed a typical oval yeast shape, whereas those in biofilms formed by strain #0390 (**Fig. 6**) mostly exhibited a more elongated (almost pseudohyphal) morphology. For all strains, inhibitory concentrations of AgNPs caused alterations in the shape and size of individual cells within the biofilms with also less cell clustering observed. In the case of *C. auris* strain #0386, low concentrations of AgNPs induced elongation of the shape in the yeast cells, similar to the pseudohyphae. However, when exposed to a higher concentration of AgNPs, the cell shape becomes spheroid, and no yeast- or pseudohyphae-shaped cells were observed (**Fig. 5**). In contrast, in the *C. auris* #0390 strain, low concentrations of AgNPs induce and enlargement of the pseudohyphae-shaped cells, growing longer than in the control (**Fig. 6**), and their presence appears to be relatively higher. However, at higher concentrations of AgNPs, the cells of this strain become yeast-shaped again but with aberrant morphology. Also, in several instances, yeast cells remained attached to each other after cell division, leading to the formation of small multi-branched chains of cells, typically in groups of less than 10 cells. Supplementary **Figure S3** includes SEM observations for the reminder of *C. auris* strains, with similar effects on biofilm structure and cellular morphology. (Supplementary **Fig S4**).

The observed effects of AgNPs on cellular morphology merit some further discussion. These effects seem to be clade-related, based on our SEM analysis for the 10 strains included in this study. Both strains from clade III were not affected in their cellular morphology. In contrast, in both strains from clade IV, inhibitory concentrations of the AgNPs altered the cell shape, leading to round-shaped cells. Interestingly, in clade I, we observed two different effects: in strains AR #0382 and #0387, the cells were altered, from the typical yeast shape to a more elongated, pseudohyphae-like shape and was not uncommon to observe mother-daughter cells attached to each other; whereas cells from strains AR #0388, 0389 and #0390 acquire an aberrant morphology and form small multi-branched chains as a result of treatment with AgNPs. In the case of *C. auris* strain AR #0381, the only one from clade II, the cells became aberrant but remained separate from each other. Other studies have shown that AgNPs disrupt the biofilm and the cell ultrastructure in other Candida species, particularly on *C albicans* (Lara et al., 2015; Vazquez-Muñoz et al., 2014; Zamperini et al., 2013), but the effect on other morphological traits, as the cell wall thickness, extracellular matrix integrity and intracellular bioaccumulation of cells remained to be further studied.

Overall our results indicate that silver nanoparticles display potent antimicrobial activity against all *C. auris* strains tested, both under planktonic and biofilm growing conditions, irrespective of their clade or geographical origin and regardless of their susceptibility and resistance patterns against clinically-used antifungals. They suggest that silver nanoparticles may be used as sanitizers and potentially in future uses to control skin colonization, and contribute to control of the nosocomial spread of this emerging pathogen.

## Abbreviations

AgNPs: Silver nanoparticles
MIC: Minimal Inhibitory Concentration
IC_50_: Inhibitory Concentration that reduces the microbial activity (XTT reading) by 50%
MFC: Minimal Fungicidal Concentration
SEM: Scanning Electron Microscopy
CDC: Centers for Disease Control and Prevention

## Acknowledgments

RV-M acknowledges the receipt of a postdoctoral scholarship from the Mexican Council of Science and Technology of Mexico (Conacyt). Support in the laboratory was provided by the Margaret Batts Tobin Foundation, San Antonio, TX, USA (to JLL-R). We thank the South Texas Center for Emerging Infectious Diseases (STCEID) for the purchase of the SEM microscope used in these studies. The funders had no role in study design, data collection, and analysis, decision to publish, or preparation of the manuscript, and the content is solely the responsibility of the authors.

## Author Contributions Statement

All authors contributed to the study design and execution, data collection and analysis, and the preparation of the manuscript. All authors have approved the final version of the manuscript.

## Contribution to the Field Statement

This study contributes to expanding the virtually non-existing knowledge regarding the effect of nanomaterials on the emergent multidrug-resistant *C. auris* strains. The antimicrobial activity of silver nanoparticles (AgNPs) was assessed in 10 *C. auris* strains with different antibiotic-resistance profiles. Additionally, the impact of AgNPs on the ultrastructure – at the cellular and the biofilm level-was assessed, which contributes to the understanding effect of AgNPs, for their potential use in the future.

