## Supplementary materials for "Silver nanoantibiotics display strong antifungal activity against the emergent multidrug-resistant yeast *Candida auris* under both planktonic and biofilm growing conditions"

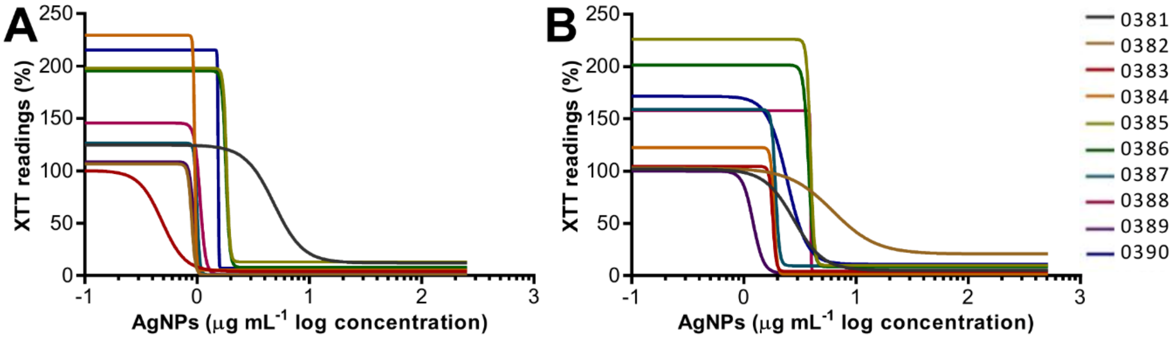


**Supplementary Figure 1. Antibiofilm activity of AgNPs against the different *C. auris* strains.** The dose-response curve shows that AgNPs display potent inhibitory activity against all the different *C. auris* strains included in this study, including inhibition of biofilm formation (panel 537 **A**) and against preformed biofilms (panel **B**).

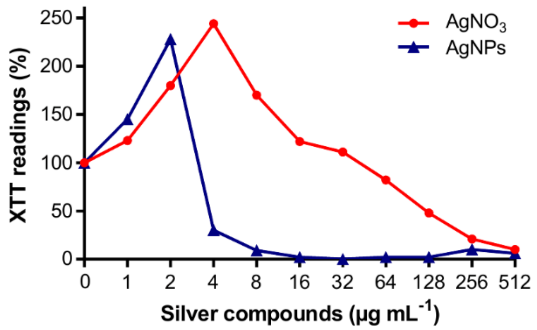


**Supplementary Figure S2. Silver compounds exhibit a paradoxical effect on *C. auris* biofilms at low concentrations**. Incubation in the presence of low subinhibitory concentrations of AgNO3 and AgNPs leads to an increase in biofilm activity (XTT readings) for biofilms formed by *C. auris* strain #0390 strain. In contrast to silver ions, this effect rapidly disappears in the case of AgNPs, turning into potent inhibitory activity at still relatively low concentrations.

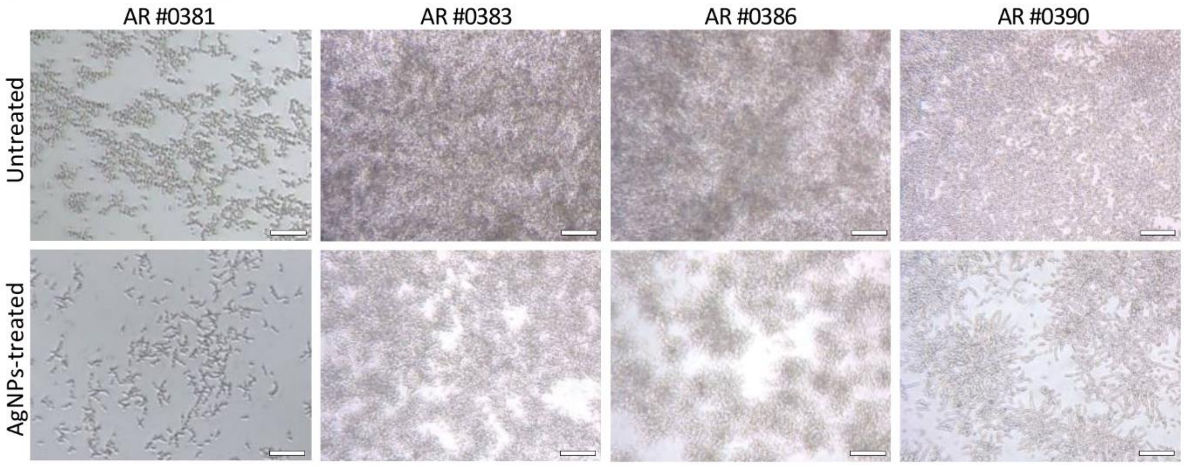


**Supplementary Figure S3. AgNPs reduce the biofilm formation in *C. auris***. Optical microscopy images reveal that subinhibitory concentrations AgNPs reduce the ability of *C. auris* to form biofilms -as seen in the reduced area of biofilm surface coverage- when compared with their respective untreated controls.

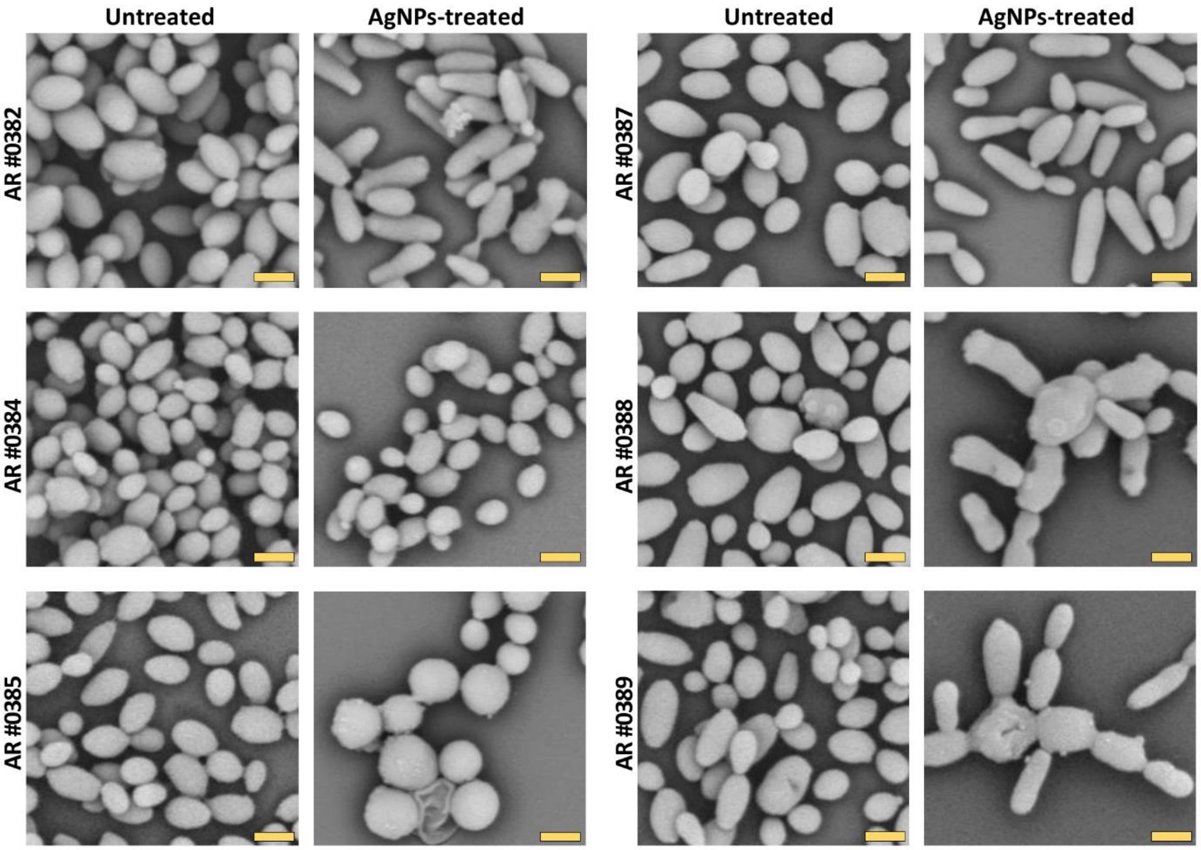


**Supplementary Figure S4. Ultrastructure analysis of the *C. auris* biofilms**. SEM images reveal that subinhibitory concentrations of AgNPs reduce the biofilm formation on all *C. auris* strains. Also, AgNPs negatively alter the shape and size of some *C. auris* strains (#0382, #0385, #0387, #0388, and #0389 strains). The effect on morphology is clade-related. Scale bar= 2 µm.
